# Follicle stimulating hormone is an autocrine regulator of the ovarian cancer metastatic niche through Notch signaling

**DOI:** 10.1101/355537

**Authors:** Sakshi Gera, Sandeep Kumar S, Shalini N. Swamy, Rahul Bhagat, Annapurna Vadaparty, Ramesh Gawari, Ramray Bhat, Rajan R Dighe

## Abstract

The association between the upregulated Notch and FSH signaling and ovarian cancer is well documented. However, their signaling has been investigated independently and only in the primary tumor tissues. The aim of this study was to investigate the interactive effects of FSH and Notch signaling on the ovarian cancer proliferation, formation and maintenance of the disseminated ovarian cancer cells. Roles of Notch and FSH in the ovarian cancer pathogenesis was investigated using ovarian cancer cell lines and specific antibodies against Notch and FSH receptor (FSHR). FSH upregulated Notch signaling and proliferation in the ovarian cancer cells. High levels of FSH were detected in the ascites of patients with serous ovarian adenocarcinoma. The spheroids from the ascites of the patients, as well as, the spheroids from the ovarian cancer cell lines under low attachment culture conditions, expressed FSHβ subunit mRNAs and secreted the hormone into the medium. In contrast, the primary ovarian tumor tissues and cell line monolayers expressed very low levels of FSHβ. The ovarian cancer cell spheroids also exhibited higher expression of the FSH receptor and Notch downstream genes than their monolayer counterparts. A combination of FSHR and Notch antagonistic antibodies significantly inhibited spheroid formation and cell proliferation *in vitro*. This study demonstrates that spheroids in ascites express and secrete FSH, which regulates cancer cell proliferation and spheroidogenesis through Notch signaling, suggesting that FSH is an autocrine regulator of cancer metastasis. Further, Notch and FSHR are potential immunotherapeutic targets for ovarian cancer treatment.

## Introduction

Ovarian cancer is the leading cause of death among the gynecological malignancies ^1^. One of the many risk factors recognized for the ovarian cancer is the excessive exposure of the normal ovarian surface epithelium to gonadotropins - Follicle stimulating hormone (FSH) and Luteinizing hormone (LH) during menopause, ovulation or infertility therapy ^2–4^. The gonadotropins are heterodimeric glycoproteins, comprised of a common α subunit associated non-covalently with the hormone-specific β subunit, secreted by the anterior pituitary and are essential for follicular development in the ovary ^5^. In addition, FSH stimulates growth of the ovarian cancer cells ^6,7^ while inhibiting apoptosis ^8,9^.

Notch signaling plays role in a wide spectrum of cell fate decisions. There are four Notch receptors (Notch1- 4) and four ligands (Jagged1 and 2, Delta 1 and 4) known in mammals. A direct link between aberrant Notch signaling and ovarian cancer progression has been previously reported ^10^.

With progression of ovarian cancer, the cells detach from the primary tumor as single cells or cellular aggregates termed as spheroid, which either remain in the ascites and contribute to the recurrence of the disease or attach to the peritoneum for the development of secondary tumors ^11^. The spheroids have been shown to be less susceptible to chemotherapy than the single cells and disruption of the spheroids re-sensitizes the ovarian tumor cells to chemotherapy with platinum-based drugs ^12,13^.

In this study, the link between FSH and Notch pathways has been investigated in more details using three different ovarian cancer cell lines. We demonstrate that FSH upregulates Notch signaling in these cell lines. Further, we demonstrate higher levels of FSH in the ascites of the ovarian cancer patients and trace the origin of this high FSH to the spheroids obtained from the patients.

## Materials and Methods

### Ovarian cancer cell lines

The ovarian cancer cell lines OVCAR-3, SKOV-3 and OVCAR-4 were authenticated by Short Tandem Repeat (STR) analysis. The OVCAR-3 cells were maintained in RPMI medium (Sigma) supplemented with 15% Fetal Bovine Serum (FBS) (GIBCO). The SKOV-3 cells were maintained in Mc-Coys medium (Sigma) and the OVCAR-4 cells were maintained in DMEM medium (Sigma) supplemented with 10% FBS. IOSE-364, a kind gift from Dr. Pritha Ray (Advanced Centre for Treatment, Research and Education in Cancer (ACTREC), Navi Mumbai) was also cultured in DMEM supplemented with 10% FBS. All media were supplemented with penicillin and streptomycin (GIBCO).

### Hormone and Antibodies

The iodination grade-purified hormones and cAMP antiserum ^39^ were obtained from the National Hormone and Pituitary Program. Polyclonal antibodies against FSH receptor (FSHR) extracellular domain (RF5) ^14, 36^ and Notch3 receptor Negative regulatory region (NRR) ^37^ were raised in the rabbits using the well-established immunization protocol ^15^. The Single chain variable fragments (ScFv) against Notch3 NRR were isolated from the yeast display library using the standardized protocol ^16^. The interesting ScFv clone (ScFv42) ^38^ was expressed in *E. coli*. *Bl21* and purified by His^6^ tag affinity chromatography.

### FSH Receptor Binding assay

Binding of FSH to the receptors present on the ovarian cancer cell lines was analyzed by radioreceptor assay. FSH was radio-iodinated using the IODOGEN method ^17^. The specific binding of ^125^I-FSH to the membrane preparations from the ovarian cancer cell lines was demonstrated as described earlier ^18^.

### *In Vitro* cAMP Measurement

Approximately 1×10^5^ OVCAR-3 cells/well were plated in a 48-well plate and 24 hours later were incubated with fresh medium containing 1mM phosphodiesterase inhibitor 3-isobutyl-1-methylxanthine (IBMX) for 30 minutes at 37°C in a CO_2_ incubator (100μ1) and then incubated with varying concentrations of FSH for 15 minutes (100μ1) after which the cells were lysed in 200μ1 of 0.2N HCl, and total cAMP produced was determined by RIA as described earlier ^14^.

### Flow cytometry-based detection of FSH receptor and Notch receptors

The ovarian cancer cells were detached from the tissue culture flasks using 0.5mM EDTA in Dulbecco’s phosphate buffer saline (DPBS) and resuspended in a medium containing 10% FBS. Approximately 1X10^5^ cells were incubated with the primary antibody (Notch3 NRR a/s or RF5 a/s) at dilution of 1:500 for 1 hour at 4°C, followed by washing thrice with DPBS and resuspension in 100μl of the medium with 10% FBS and 1:1000 dilution of anti-rabbit IgG-FITC (Invitrogen) for 45 minutes at 4°C. After incubation, the cells were washed and resuspended in DPBS and analyzed using the Becton Dickinson Accuri and the median fluorescence values were analyzed.

### Notch signaling assays

Notch signaling in the ovarian cancer cell lines was determined as described earlier ^19^. Briefly, the ovarian cancer cell lines were seeded (5X10^4^ cells/well) in 24 well plates (Nunc) and transfected with 790ng of 12XCSL (CBF1/Su(H)/Lag-1) reporter plasmid and 10ng of pGL3 basic or 800ng of pGL3 control along with 1ng of pRL-Tk using Lipofectamine 2000 (Invitrogen) as per the manufacturer’s instructions. The 12XCSL reporter vector was a kind gift from Prof Urban Lendahl ^20^. The vector contains the multimerized high affinity CSL sites upstream of luciferase reporter gene. The Notch Intra cellular domains after activation, translocate to the nucleus and acts as a transcriptional regulator of CSL dependent pathways and thus upregulates the expression of luciferase reporter in transfected cells. The transfected cells were incubated with increasing concentrations of FSH in presence or absence of RF5 a/s, Notch inhibitor N-[N-(3,5-difluorophenacetyl)-l-alanyl]-S-phenyl-glycine t-butyl ester (DAPT), 5μM (Sigma) or ScFv42 (10μg/ml) and the luciferase reporter activity was estimated using the Dual-luciferase assay kit (Promega) in TD200-Luminometer.

### Ovarian cancer cell line proliferation

Proliferation of the ovarian cancer cell lines was determined using 5-bromo-2’-deoxyuridine (BrdU) incorporation assay. Briefly, 5 × 10^3^ cells were seeded per well in a 96-well plate and synchronized by incubating in medium without FBS overnight. Next day, the cells were treated with FSH or FSHR and Notch antagonists in the medium supplemented with FBS. The cells were labeled with BrdU for 12 hours and its incorporation was determined as per manufacturer’s recommendation (Calbiochem, USA).

### Expression analysis by Real time PCR

Total RNA from the ovarian tumors or spheroids obtained from the patients’ peritoneal tap, or from the ovarian cancer cell lines was isolated by Trizol-based method. Briefly, the cells were washed with DPBS and lysed in Tri-reagent (Sigma). RNA was extracted according to the manufacturer’s recommended protocol and cDNA was synthesized from 1μg of total RNA using the cDNA preparation kit (Thermo Scientific) using the random primers. The transcript levels of the interested genes were analyzed using the SYBR-Green in Eppendorf Realplex thermo-cycler. Glyceraldehyde 3-phosphate dehydrogenase (GAPDH). The ΔΔCt method was used to calculate the fold difference in transcript levels of different genes. The absolute expression of FSH and FSHR in primary tumors and spheroids was intrapolated from the dilution curve of the respective genes generated in the same experiment. The sequences of the primers used in the analysis listed in the Table 1.

**Table 1.**
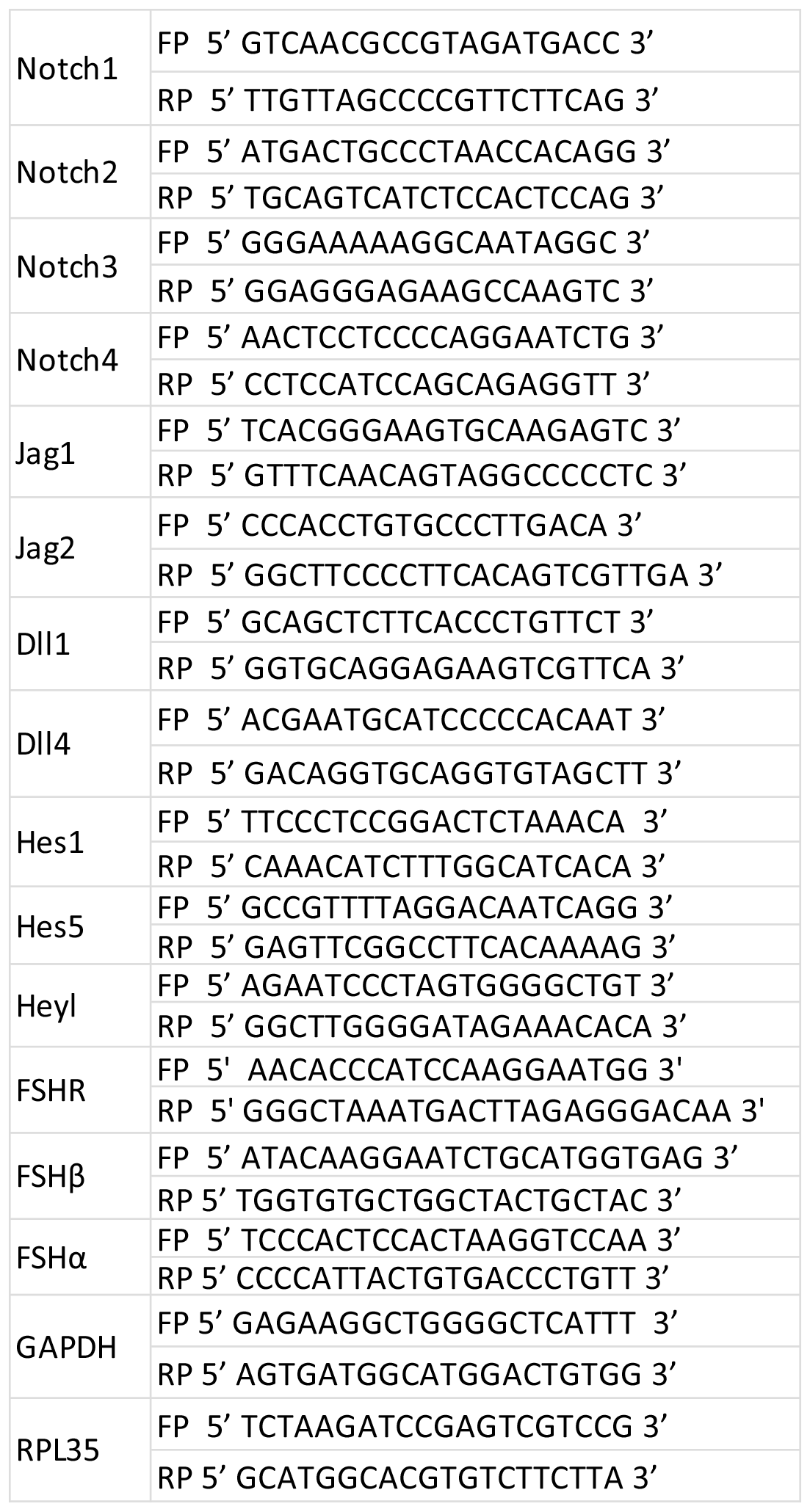
List of real time primers.

### Estimation of FSH in patients’ ascites

The FSH levels in the ascites and sera of the patients as well as the hormone secreted by the ovarian cancer cell lines were determined by RIA ^21^.

### Spheroid formation by ovarian cancer cell lines

Cellular aggregates of the ovarian cancer cell lines were developed by culturing the cells under low attachment conditions. Briefly, the cells (~10^4^/well) were seeded in 96 well low attachment plates (NUNC sphera) and allowed to form aggregates for 24 to 96 hours. The aggregates were treated with RF5 a/s and/or ScFv42 at different time points. At regular intervals, images of the aggregates were captured using the Zeiss microscope at 60X zoom. Effect of different treatments on cellular aggregation or disaggregation was determined by analyzing the area of aggregates in a script developed in MATLAB-R 15a (100 pixels in the image was equivalent to 150μM) fixing the ellipticity constant at 0.1.

### Statistical analyses

The statistical analyses were performed using the Graphpad Prism 5.0 software. The p-value of < 0.05 was considered statistically significant. Each experiment was repeated three times.

## Results

### FSH and Notch signaling in the ovarian cancer cell lines

Presence of the functional FSHR and Notch receptors in the ovarian cancer cell lines (OVCAR-3, SKOV-3 and OVCAR-4) was demonstrated using different approaches. Presence of FSHR in these cells was demonstrated by determining specific binding of ^125^I-FSH to the ovarian cancer cell membranes. The membrane preparations were incubated with ^125^I FSH in presence of increasing concentrations of unlabeled FSH and the binding data obtained were converted into Scatchard plots. As shown in Figure 1A, the dissociation constants of the OVCAR-3 and SKOV-3 were 1.7×10^−9^M and 1.9×10^−9^M respectively, while that of the OVCAR-4 was 2.8×10^−9^M. The B_max_ for the receptor was highest in the OVCAR-3 cells (Figure1A).

**Figure 1.**
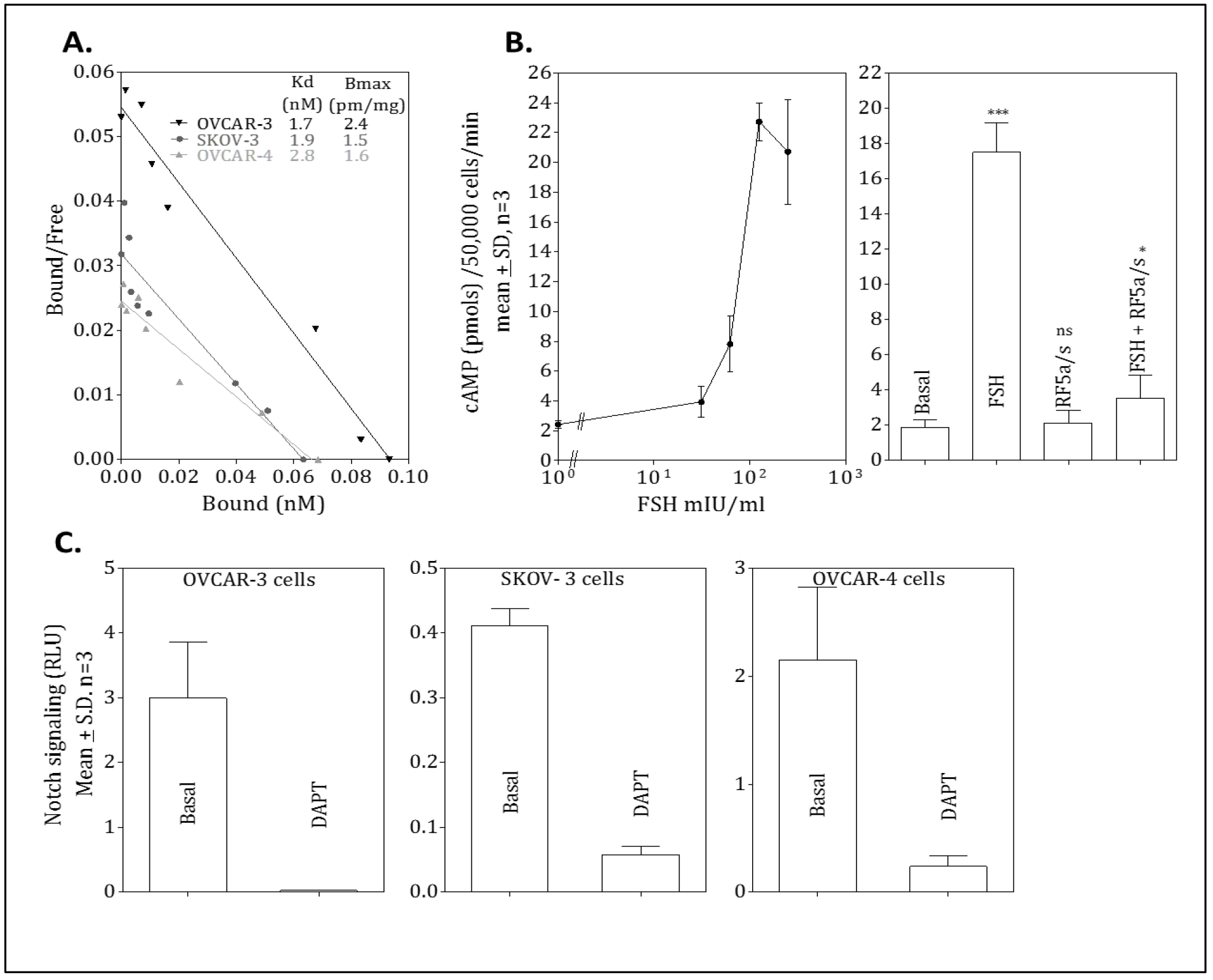
Characterization of FSH signaling in ovarian cancer cell lines. **A.** The presence of FSHR on the ovarian cancer cell lines OVCAR-3, SKOV-3 and OVCAR-4 cell was investigated by determining the binding of ^125^I-FSH to their membranes (10μg each) in the presence of increasing concentrations of cold FSH. The binding data were converted into a Scatchard plot and the inset table shows the K_d_ and B_max_ of hormone in the three cell lines. **B.** The activity of FSH receptor was investigated by analyzing the levels of cAMP in the presence of hormone by radioimmunoassay (RIA). The OVCAR-3 cells were incubated with increasing concentrations of FSH or **C.** with 100mIU/ml FSH in the presence and absence of 5μg/ml of RF5 a/s and cAMP levels were determined. Significance is calculated by comparing the treated with basal using non-parametric unpaired T test where ns represents P> 0.05, * represents P≤ 0.05, and *** represents P≤ 0.001. Error bars represent mean ± Standard Deviation (S.D.) and n is the number of repeats. **D.** The ovarian cancer cells were transfected with 12XCSL luciferase reporter plasmid and incubated with 5μM DAPT for determining the Notch signaling activity. The reporter activity was analyzed after 24 hours of incubation by dual luciferase assay. RLU, Relative Luciferase Unit.

Presence of FSHR on the ovarian cancer cells was also confirmed by flow cytometry using the antiserum raised against FSHR (RF5 a/s) ^14^. The cells were incubated with RF5 a/s and its binding was determined using anti-Rabbit IgG-FITC. As shown in the Figure 2A, all three ovarian cancer cell lines exhibited binding of the a/s indicating the presence of FSHR on their surfaces with the OVCAR-3 cells displaying highest antibody binding, which agrees with the Scatchard analysis. Further, six-fold higher expression of FSH receptor was observed at mRNA level in OVCAR-3 cells compared to normal ovarian epithelial cell line IOSE 364.

**Figure 2.**
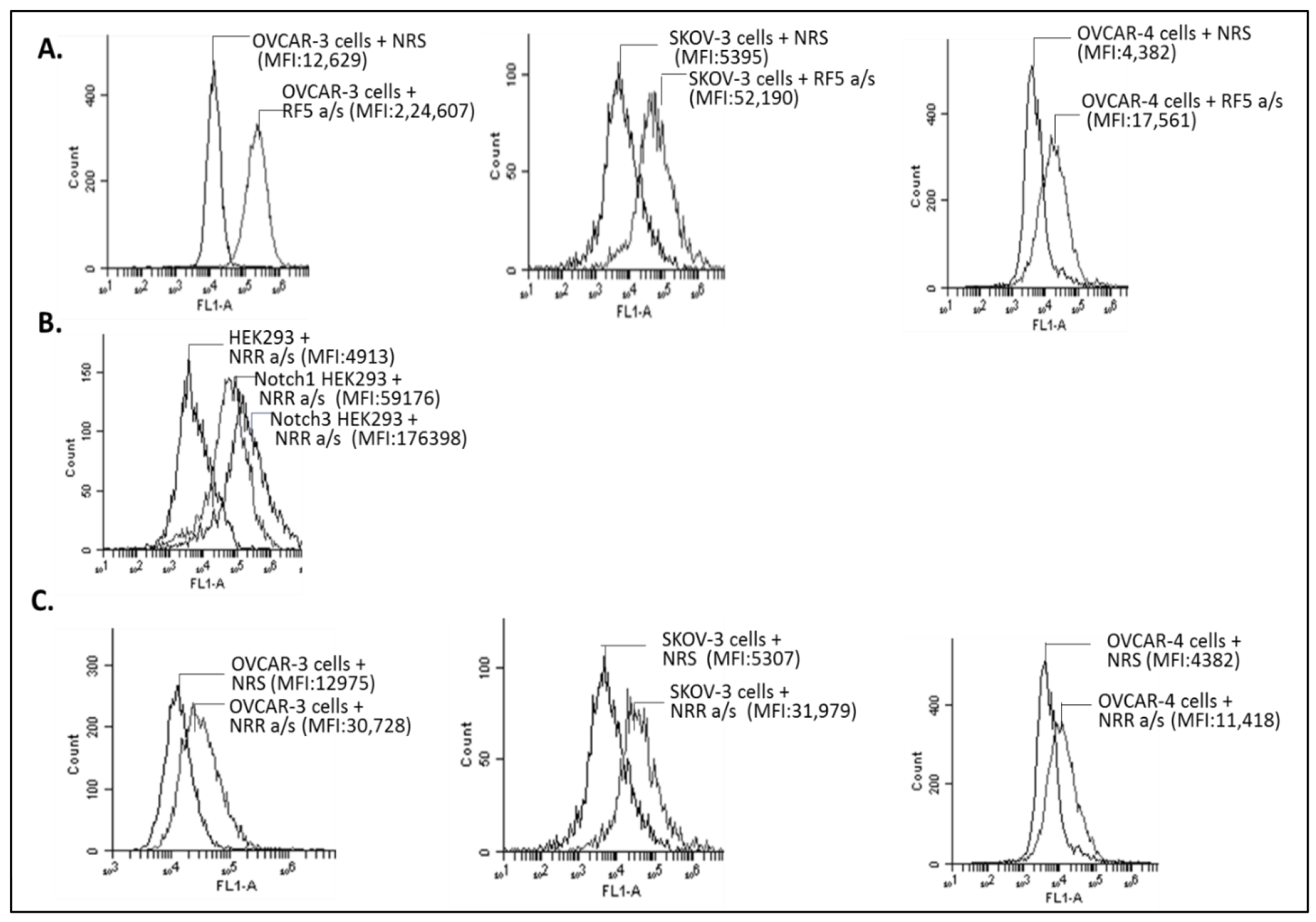
FSHR on ovarian cancer cell lines. The presence of FSHR on the ovarian cancer cell lines OVCAR-3, SKOV-3, and OVCAR-4 was analyzed by flow cytometry. The ovarian cancer cell lines were incubated withRF5 a/s (1:500 dilution) or Normal rabbit a/s (NRS) and analyzed for binding using FITC-conjugated anti-rabbit secondary antibody. The histograms are representive of three independent experiments. **Notch receptors on ovarian cancer cell lines. B.** HEK293 cells and Notch1 and Notch3 overexpressing HEK293 cells were incubated with Notch3 NRR a/s (1:500 dilution) followed by anti-rabbit FITC and binding was analyzed by flow cytometry. **C.** The presence of Notch receptor on the ovarian cancer cell lines OVCAR-3, SKOV-3 and OVCAR-4 was analyzed by flow cytometry. The ovarian cancer cell lines were incubated with Notch3 NRR a/s (1:500 dilution) or Normal rabbit a/s (NRS)and analyzed for binding by antirabbit secondary antibody conjugated with FITC. The histograms are represented of three independent experiments (MFI, Median fluorescent intensity, FITC, Fluorescein Isothiocyanate).

The functional nature of FSHR was further confirmed by determining response of the OVCAR-3 cells to FSH *in vitro*. The cells were incubated with increasing concentrations of FSH for 15 minutes at 37°C and cAMP formed was determined by RIA. As shown in Figure 1B, FSH produced a dose-dependent increase in cAMP. In the next experiment, the OVCAR-3 cells were incubated with FSH in presence and absence of RF5 a/s, and cAMP produced was determined. As shown in Figure 1B, response to the hormone was significantly inhibited in presence of RF5 a/s indicating the antagonistic nature of the a/s ^14^.

Presence of Notch receptors on the ovarian cancer cell lines was demonstrated by flow cytometry using the antiserum raised against the NRR of Notch3 that has full cross reactivity with the Notch1 (Figure 2B). As shown in Figure 2C, all three ovarian cancer cell lines exhibited binding of the NRR a/s indicating presence of Notch receptors on their cell surface. Notch signaling in the OVCAR-3 cells, as determined by 12X CSL reporter, was stimulated by Dll4 (see below, Figure 3C) and inhibited by DAPT (Figure 1C).

**Figure 3.**
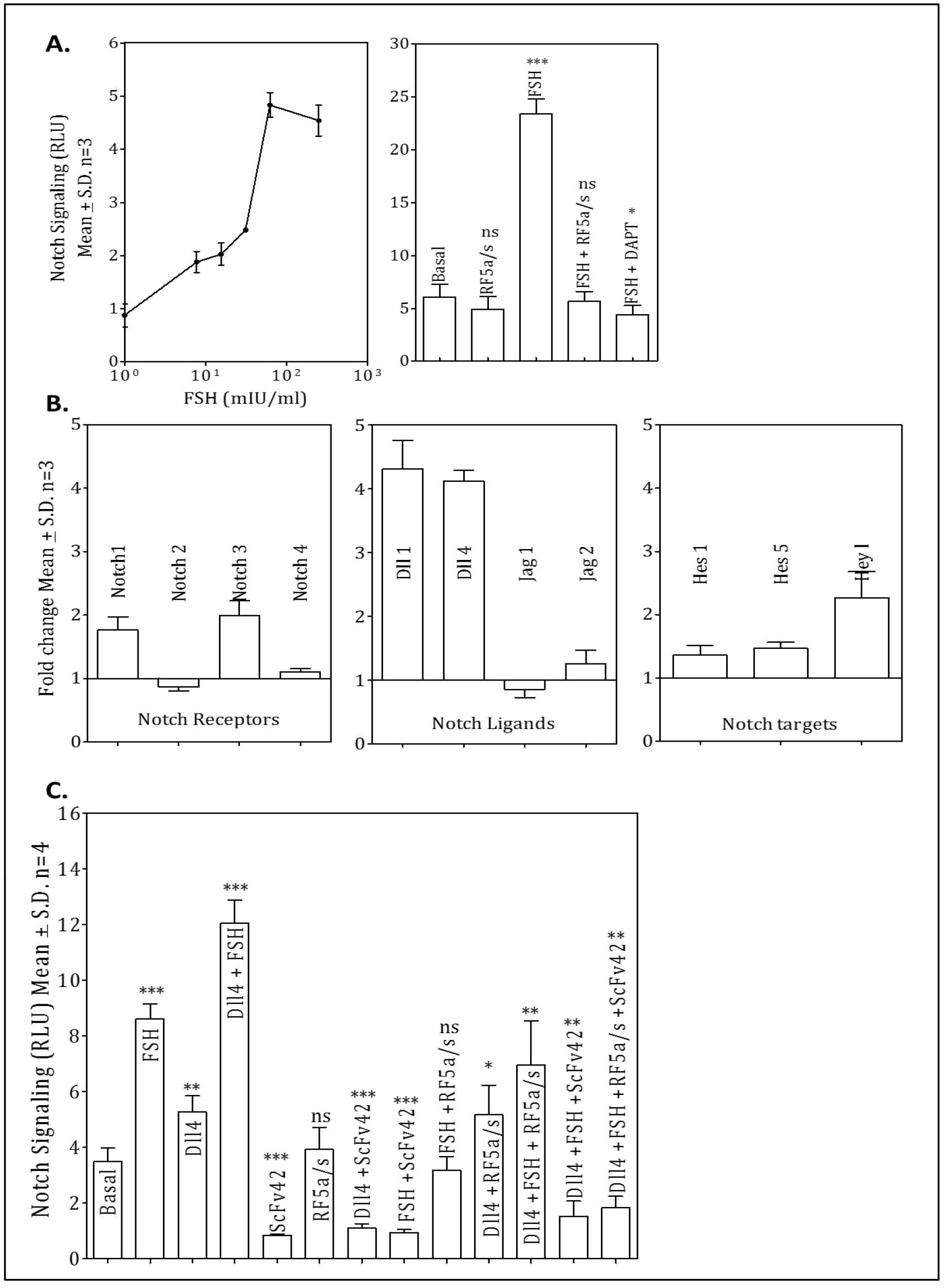
Effect of FSH on Notch signaling. **A.** The OVCAR-3 cells were transfected with 12XCSL luciferase reporter plasmid and incubated with increasing concentrations of FSH or 100mIU/ml of FSH in the presence of 5μg/ml FSHR antagonist RF5 a/s and 5μM DAPT in different combinations for 48 hours. The reporter activity was measured by dual luciferase assay after 48 hours of incubation. **B**. The OVCAR-3 cells were incubated with 100mIU/ml of FSH for48 hours and real time PCR was performed for indicated Notch receptors, ligands and target genes. The fold change in the expression of receptor and ligands was calculated with respect to cells cultured without hormone after normalizing with GAPDH expression. **C.** OVCAR-3 transfected with 12XCSL luciferase reporter plasmid were cultured on pre-coated Dll4-Fc. 100mIU/ml of FSH, or 5μg/ml FSHR antagonist RF5 a/s, or 10μg/ml ScFv42, or in different combinations were added to the cells for 48 hours and the effect on the Notch signaling was measured by dual luciferase assay. RLU is Relative luciferase unit. Significance is calculated by comparison of treated and basal values using non-parametric unpaired T test where ns represents P> 0.05, * represents P≤ 0.05, ** represents P≤ 0.01 and *** represents P≤ 0.001 compared to basal values. Error bars represent mean ± SD and n is the number of repeats.

### Effect of FSH on Notch signaling in the ovarian cancer cell lines

Possible effect of FSH on Notch signaling in the ovarian cancer cell lines was investigated by incubating the cell lines with increasing concentrations of FSH for 48 hours and determining Notch signaling in cells expressing 12XCSL luciferase reporter. As shown in the Figure 3A, in OVCAR-3 cells, there was FSH dose-dependent increase in luciferase activity indicating enhanced Notch signaling, which was further confirmed by increase in the transcript levels of Notch target genes Hes1, Hes5 and Heyl (Figure 3B). Similar results were obtained for SKOV-3 and OVCAR-4 cells (Figure 4A and B). The stimulatory effect of FSH on Notch signaling was completely inhibited by the RF5 a/s, which by itself had no effect on Notch signaling. The hormone stimulated Notch signaling was also inhibited by DAPT (Figure 3A).

**Figure 4.**
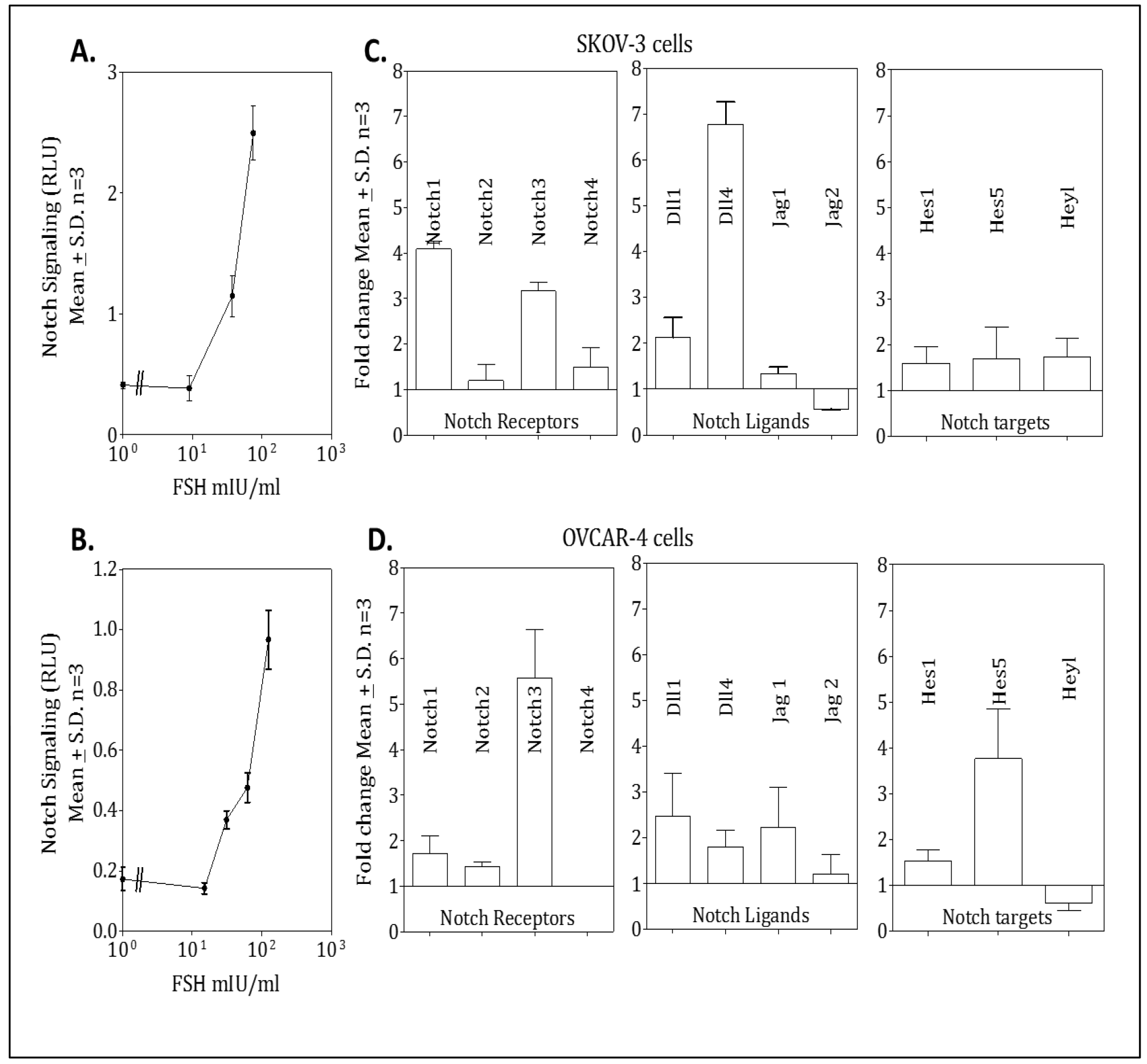
Effect of FSH on Notch signaling. **A.** SKOV-3, and **B.** OVCAR-4 cells were transfected with 12XCSL luciferase reporter plasmid and incubated with increasing concentrations of FSH. The effect on the Notch signaling was measured by dual luciferase assay after 48 hours. RLU, Relative luciferase unit. Error bars represent mean ± SD. n is the number of repeats. **C.** SKOV-3 and **D.** OVCAR-4 cells were incubated with 100mIU/ml of FSH for 48 hours and real time PCR was performed for indicated Notch receptors, ligands and target genes. The fold change in the transcript levels of Notch genes was calculated with respect to the levels in cells cultured without hormone after normalizing with GAPDH expression. Error bars represent mean ± SD. n is the number of repeats.

FSH upregulated expression of Notch1 and Notch3, as well as, the Notch ligands Dll1 and Dll4 in OVCAR-3 cells as determined by RT-PCR (Figure 3B). In addition, increase in Notch receptors and Dll4 was also demonstrated by flow cytometry (Figure 5). In case of OVCAR-4 cell line, upregulation was observed predominantly in case of Notch3 and the three ligands, Jag1, Dll1 and Dll4 (Figure 4B). Thus, increased Notch signaling in the ovarian cancer cell lines in response to FSH was a result of an increased expression of both Notch receptors and ligands.

**Figure 5.**
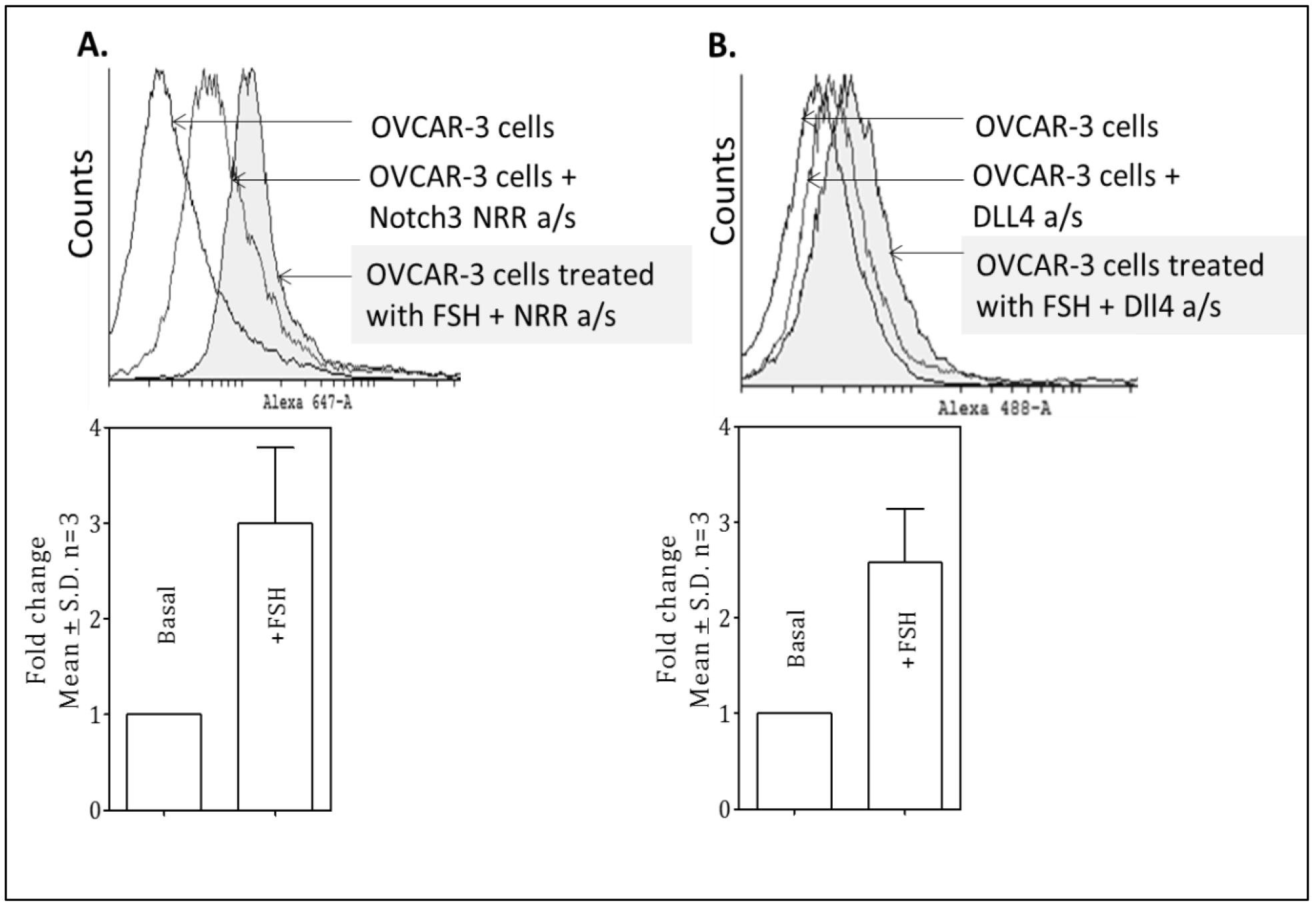
Effect of FSH on Notch receptors and ligands. The OVCAR-3 cells were harvested with EDTA after incubation with 100mIU/ml of FSH for 48 hours and labelled with **A.** Notch3 NRR a/s and **B.** Dll4 antibody for analysis by flow cytometry. The antibody bound was determined by anti-rabbit Alexa 647 and anti-mouse Alexa 488 respectively. Binding of a/s was analyzed by comparing the median fluorescent intensity in the presence and absence of hormone. The histograms are representative of three independent experiments. Quantitative analysis of the same is represented below. Significance is calculated by comparison of treated and basal values using non-parametric unpaired T test. Error bars represent mean ± SD. n is the number of repeats.

Dll4 present on the surface of the stromal cells in the ovarian cancer microenvironment is involved in progression of the ovarian tumor ^22^ and as shown above, FSH upregulates Dll4 expression. Therefore, effect of exogenous Dll4 ^19^ together with FSH on the Notch signaling was determined. The OVCAR-3 cells were incubated with FSH (100 mIU/ml) and the immobilized Dll4, individually and in combination, and Notch signaling was determined. As shown in Figure 3C, both ligands stimulated Notch signaling in an additive manner. Effect of RF5 a/s and Notch inhibitory ScFv42 on Notch signaling was investigated in the same experiment. ScFv42 is a Notch3 NRR specific ScFv and can inhibit the basal and FSH stimulated Notch signaling in a dose dependent manner (Figure 6). As shown in Figure 3C, RF5 a/s inhibited FSH-stimulated Notch signaling while ScFv42 inhibited the basal, Dll4-, FSH- and FSH+Dll4-stimulated Notch signaling. A combination of RF5 a/s and ScFv42 inhibited Notch signaling to the same extent as the ScFv42 alone.

**Figure 6.**
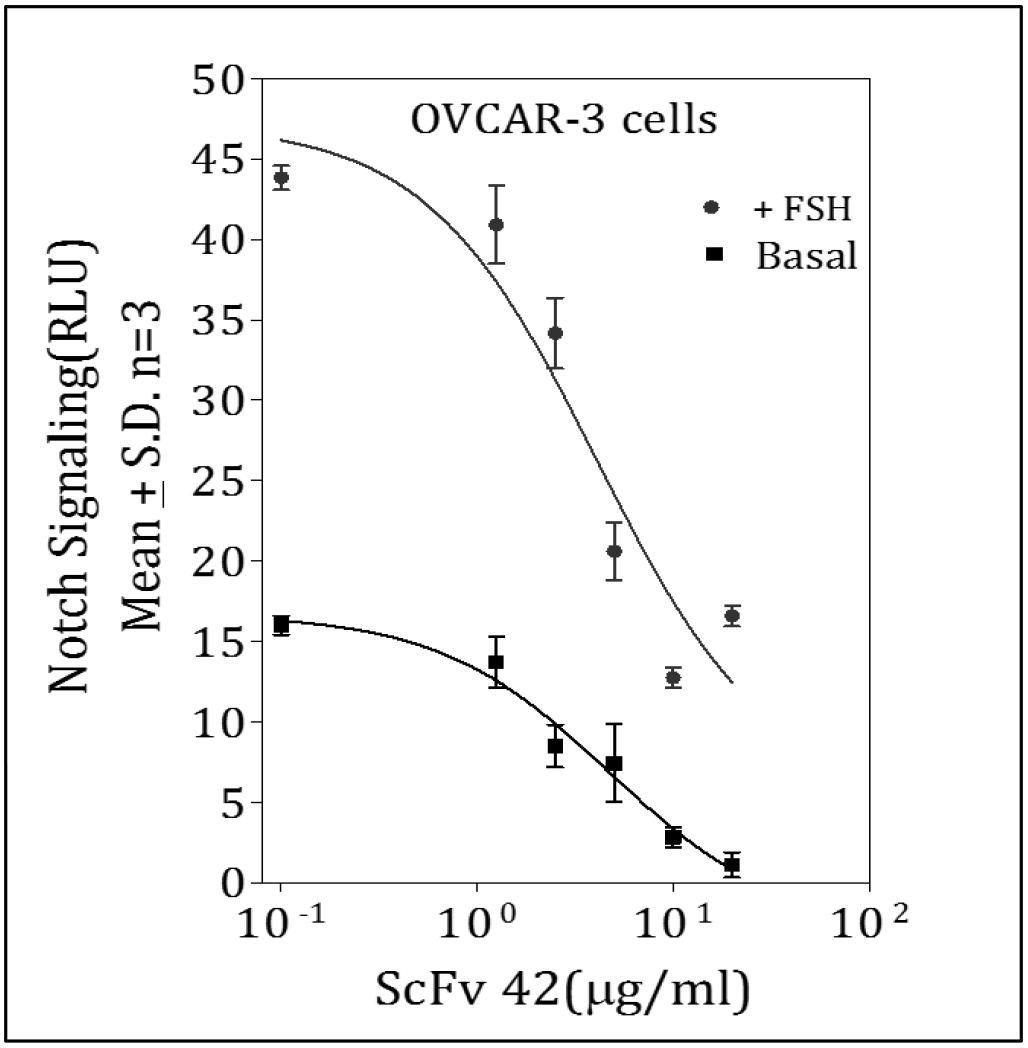
Effect of anti Notch3 ScFv on Notch signaling. OVCAR-3 cells were transfected with luciferase plasmid and incubated with increasing concentrations of ScFv42 (Notch3 NRR antagonist) in the presence or absence of 100mIU/ml FSH. The reporter activity was measured by dual luciferase assay after 48 hours of incubation. RLU, Relative luciferase unit. n is the number of repeats.

### Effect of FSH and Notch ligand on the proliferation of ovarian cancer cell lines

Role of Notch signaling in proliferation of ovarian cancer cell line is well documented while as shown above, FSH upregulated Notch signaling in the ovarian cancer cell lines. Therefore, effect of FSH on proliferation of OVCAR-3 cells was determined. The OVCAR-3 cells were incubated with increasing concentrations of hormone for 48 hours and BrdU incorporation was analyzed. As shown in Figure 7A, FSH stimulated OVCAR-3 cell proliferation in a dose-dependent manner.

**Figure 7.**
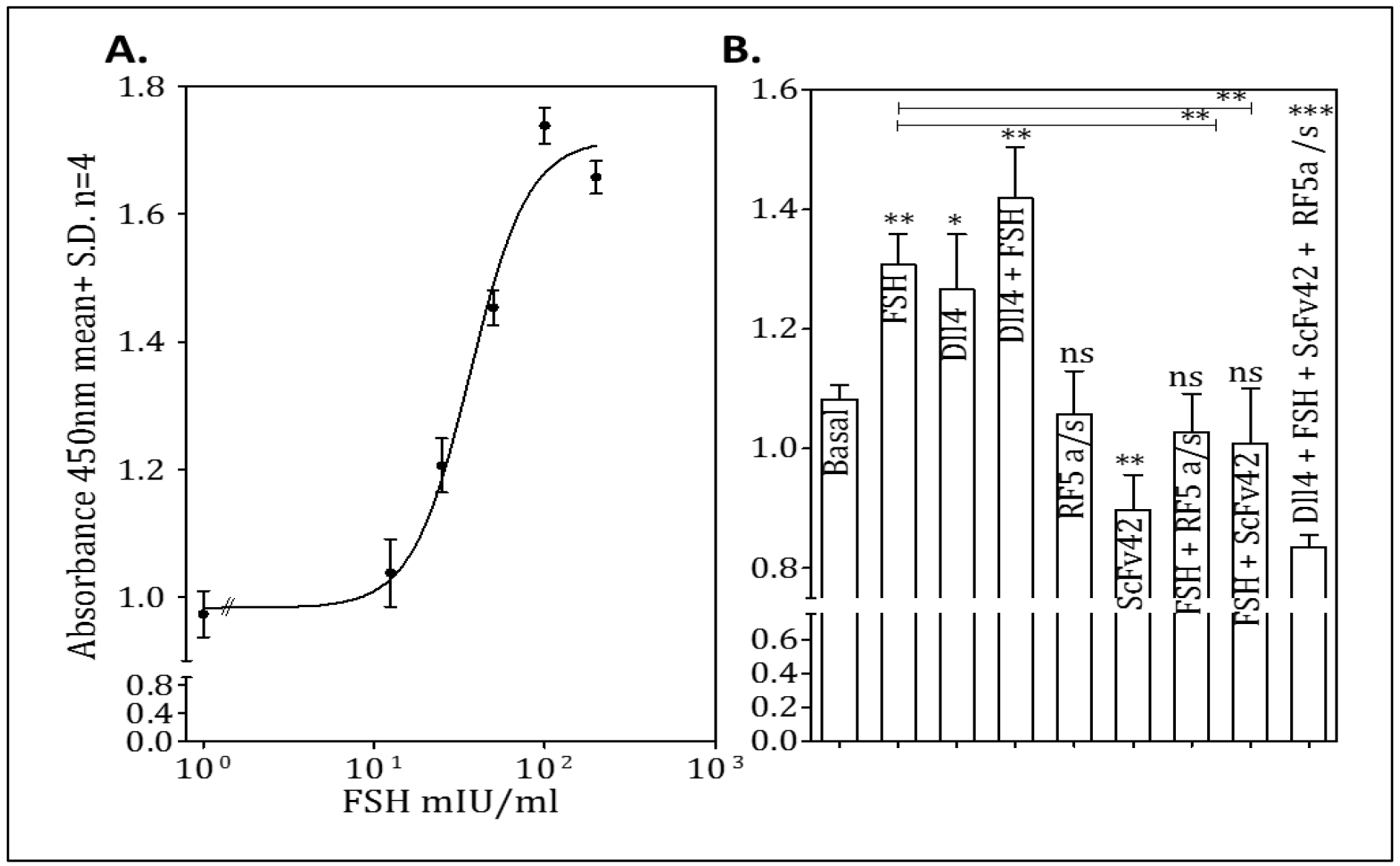
Effect of FSH on proliferation of ovarian cancer cells. **A.** The OVCAR-3 cells were synchronized by culturing them in serum free medium overnight and incubated with increasing FSH concentrations. The proliferation rate was investigated by analyzing the BrdU incorporation in the cells after 48 hours. **B**. OVCAR-3 cells (synchronized as above) were cultured on pre-coated Dll4-Fc and incubated with 100mIU/ml of FSH, or 5μg/ml FSHR antagonist RF5 a/s, or 10μg/ml ScFv42, or in different combinations for 48 hours. Proliferation rate was investigated by BrdU incorporation and significance was calculated by unpaired T test compared to the basal levels where ns represents P> 0.05, * represents P≤ 0.05, ** represents P≤ 0.01, *** represents P≤ 0.001 compared to basal. n is the number of repeats. Error bars represent mean ± SD.

Next, effects of FSH and Notch ligand, individually and in combination, on proliferation of OVCAR-3 cell line were examined. As shown in Figure 7B, both ligands stimulated cell proliferation in an additive manner. This increase in proliferation was inhibited by RF5 a/s, as well as, ScFv42. Further, the FSH- and Dll4-stimulated proliferation was inhibited by both RF5 a/s, as well as, the Notch3 NRR-specific ScFv even below the basal level.

### FSH levels in ovarian cancer patients’ tumors, sera and ascites

To understand the role of FSH and Notch signaling in the ovarian cancer progression, FSH levels in the ascites of cancer patients were estimated. Ascites samples were obtained by peritoneal tap from 26 ovarian cancer patients while serum samples were obtained from seven patients and FSH levels were determined by RIA. FSH was detected in ascites of all patients with levels ranging from 15 mIU/ml to 208 mIU/ml. The serum could be the possible source of FSH in the ascites. Average serum FSH levels in the post-menopausal Indian women (age 54 ± 8 years) were 62 ± 30 and 25.8±10 mIU/ml in the premenopausal women ^23,24^ As shown in the Table 2, the ascitic FSH levels in twelve patients (# 2, 3, 6, 10, 11, 12, 13, 14, 17, 18, 23, 25) were more than 62 mIU/ml while the seven patients who had low FSH in their ascites, either did not have malignant cells detected in their ascites (# 1, 8, 9, 15) or were undergoing chemotherapy (# 20, 21, 24). Of the seven patients, whose serum samples were available for analysis, three patients (# 23, 25, 26) showed high FSH levels in both sera, as well as, ascitic fluids (> 100 mIU/ml). However, in the case of remaining 4 patients, there was no correlation between the serum and ascitic FSH levels.

**Table 2.** FSH in patients’ ascites and serum. Details of the patients from whom ascites, serum and ovarian tumor samples were obtained. Levels of FSH in the ascites and serum obtained from the ovarian cancer patients were determined by Radioimmune assay (RIA) in a single experiment.

| Pt # | Age (yrs) | Description | Ascitic FSH (mIU/ml) | Serum FSH (mIU/ml) |
| --- | --- | --- | --- | --- |
| 1 | NA | Reactive mesothelial cells in a hemorrhagic background with a few inflammatory cells | 22.2 | NA |
| 2 | 70 | High Grade adenocarcinoma of ovary | 71.04 | NA |
| 3 | 53 | Metastatic adenocarcinoma of ovary, | 124.14 | NA |
| 4 | 58 | High grade serous papillary carcinoma, | 45.2 | NA |
| 5 | 50 | High grade serous adenocarcinoma, Recurrent disease, stage IVB | 38 | NA |
| 6 | 72 | Serous adenocarcinoma | 102.8 | NA |
| 7 | 49 | Adenocarcinoma of ovary | 36.6 | NA |
| 8 | 40 | Shows normal squamous cells no abnormal cells were seen in cytology. | 34.2 | NA |
| 9 | 56 | Peritoneal fluid shows reactive mesothelial cells. No malignant cells seen in ascites | 13.5 | NA |
| 10 | 42 | Serous adenocarcinoma | 136.32 | NA |
| 11 | 50 | Serous papillary adenocarcinoma | 134.24 | NA |
| 12 | 48 | NA | 98.24 | NA |
| 13 | 45 | Serous adenocarcinoma | 90.96 | NA |
| 14 | 55 | High grade serous carcinoma. Had completed | 96.32 | NA |
| 15 | 68 | High grade serous papillary carcinoma. Patient undergoing CT at sample collection | 15 | NA |
| 16 | NA | NA | 40.9 | NA |
| 17 | NA | Serous adenocarcinoma, 5 cycles of CT completed | 102.77 | NA |
| 18 | 54 | High grade serous papillary carcinoma | 104.85 | NA |
| 19 | 53 | High grade serous papillary carcinoma | 35.01 | NA |
| 20 | 55 | Post NACT, 3 cycles of CT | 23.41 | 24.44 |
| 21 | 43 | No residual disease | 24 | 84.55 |
| 22 | 64 | Serous adenocarcinoma, Stage III C | 40.83 | Nil |
| 23 | 58 | Serous adenocarcinoma, Grade 1 | 208.32 | 122.19 |
| 24 | 46 | Post NACT, Stage III, Serous adenocarcinoma | 14.76 | 5.85 |
| 25 | 55 | Serous adenocarcinoma | 115.44 | 131.43 |
| 26 | 73 | Post NACT, 3 cycles of CTs, Serous papillary | 59.71 | 147.26 |
| 27 | 56 | Metastatic adenocarcinoma of ovary | NA | NA |
| 28 | 61 | Mucinous low grade ovarian carcinoma | NA | NA |
| 29 | 46 | low grade ovarian carcinoma | NA | NA |
| 30 | 59 | Ovarian malignant neoplasm | NA | NA |
| 31 | 73 | High grade serous carcinoma. Undergoing CT, post NACT | NA | NA |
| NA: Not available, CT: chemotherapy, NACT: Neoadjuvant chemotherapy |  |  |  |  |

To identify the source of the hormone in the ascites, the primary ovarian tumor tissues from 5 ovarian cancer patients (Patient # 27-31) were obtained. The total RNA was isolated and subjected to RT-PCR analysis for FSHβ, FSHα and FSHR transcripts for the absolute quantification of RNA levels. The expression was quantified based on a dilution curve of known DNA concentrations for each gene. In addition, as a comparative positive control, human pituitary tissue was included for the expression of genes encoding FSHα, FSHβ and FSH receptor. As shown in the Figure 8, patients’ tumor samples exhibited lower expression of FSH α and β suggesting very low hormone message in the tumors whereas the FSHR transcript levels were very high (Patient # 27, 28, 29 and 30). The other possibility of FSH being expressed by the metastatic tumor epithelium represented by spheroids in the ascitic fluids was next explored. Multicellular spheroids from the ascitic fluid of 8 patients (Patient # 2, 3, 5, 10, 11, 13, 14 and 18) were isolated from the cell mass obtained from the ascites from the patients and as shown in the Figure 8, the spheroids obtained from the patients 2, 3, 10, 11, 13, 14 and 18 exhibited high levels of FSHβ message except from those obtained from the patient 5, who had recurrent disease (Table 2). The FSHβ transcript levels were consistent with the ascitic hormone levels. Interestingly, there was a significantly high expression of FSHR in the spheroids of the ovarian cancer patients (2, 3, 10, 11, 13, 14 and 18) compared to the patient 5 (Figure 8).

**Figure 8.**
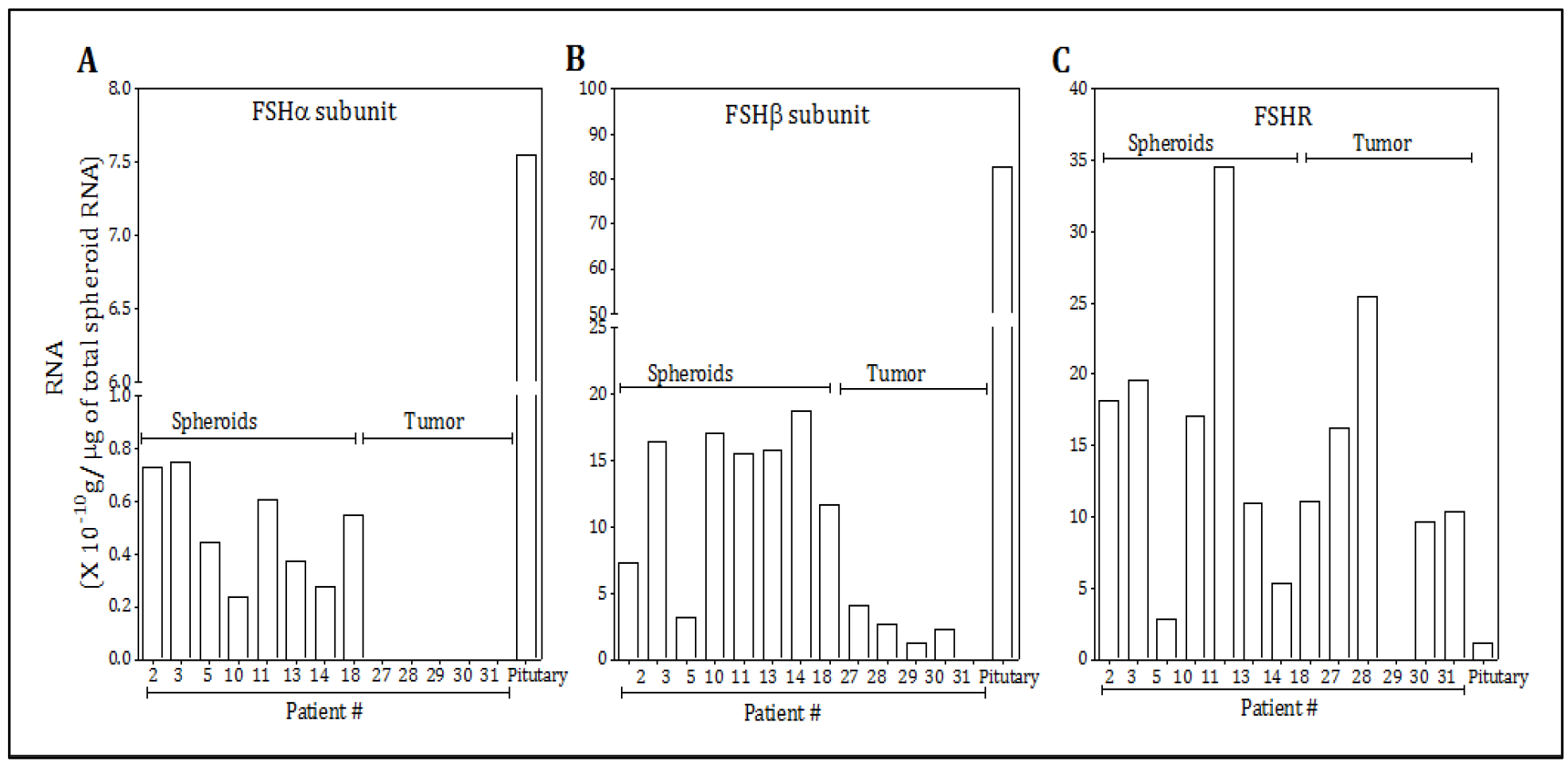
Expression of FSH and FSHR in primary tumor and spheroids obtained from ovarian cancer patients. Total RNA was isolated from the primary tumors and spheroids obtained from the ascites of ovarian cancer patients and cDNA was prepared from 1 μg of total RNA. The Cycle threshold values (Ct) of **A.** FSH α subunit, **B.** β subunit and **C.** FSHR were examined by real time PCR and the RNA levels were intrapolated from the dilution curves of the respective genes.

### Expression of FSH in ovarian cancer cell line spheroids and monolayers

To simulate the conditions of *in vivo* spheroid formation, the ovarian cancer cell lines were cultured under the low attachment conditions that facilitated the cells to aggregate and form compact spheroidal aggregates ^25^. As shown in the Figure 9A, spheroid-like aggregates of the OVCAR-3 cells were observed after 48 hours of culture. The SKOV-3 and OVCAR-4 cells formed aggregates of different geometries under these conditions (Figure 9B and 9C). Expression of FSH by the spheroids/aggregates was demonstrated by determining the hormone levels in the culture supernatants collected after different intervals and FSHβ transcript levels by RT-PCR. As shown in Figure 9D and 9E, there was an increase in FSHβ transcripts under non-attachment conditions with corresponding secretion of FSH by OVCAR-3 and SKOV-3 cells, while there was neither any increase in the FSHβ transcripts nor in secreted FSH when the cells were grown as monolayer cells. The OVCAR-4 cells did not show any increase in FSHβ transcripts, as well as, the secreted hormone (Figure 9F).

**Figure 9.**
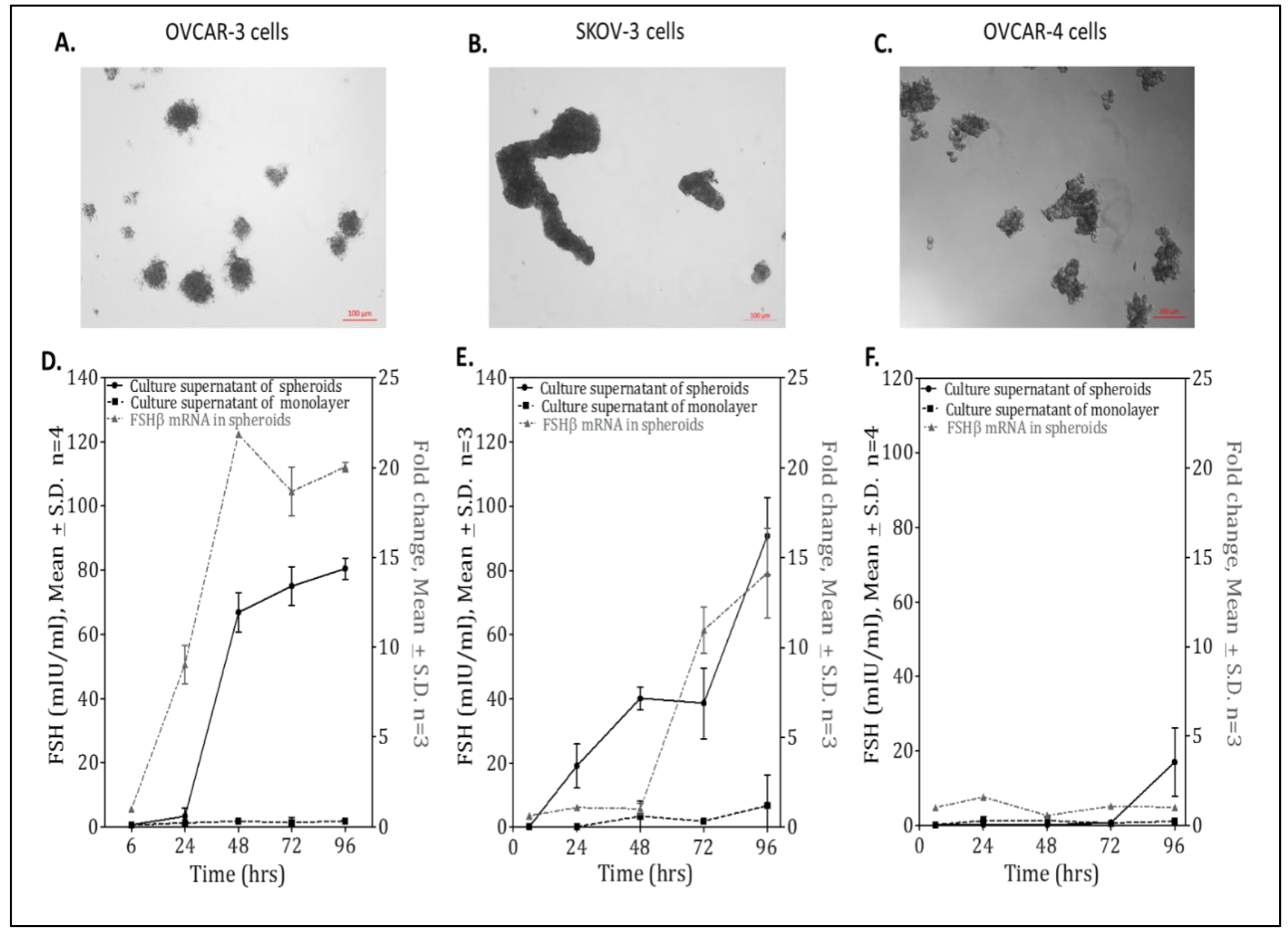
Expression of FSH in ovarian cancer cell lines spheroids. Photomicrographs images of cellular aggregates of **A.** OVCAR-3, **B**. SKOV-3 and **C.** OVCAR-4 cells cultured in low attachment conditions. **D.** OVCAR-3, **E**. SKOV-3 and **F.** OVCAR-4 cultured in low attachment conditions or as monolayer. FSH level in the conditioned media of aggregates/ spheroids or monolayers cultured for different time periods were analyzed by RIA. Real time-PCR was performed to assay the transcript levels of FSHβ in cellular aggregates. Fold change was calculated with respect to cells cultured as monolayers after normalizing with GAPDH expression. Significance is calculated by comparison of treated and basal values using non-parametric unpaired T test. n is the number of repeats. Error bars represent mean ± SD.

### Effect of FSH and Notch inhibitors on OVCAR-3 spheroids

Formation of ovarian cancer cell line spheroids mimics the ovarian tumor metastatic condition and these spheroids exhibit chemo-resistance as they increase in size^26^. Higher chemo-resistance has also been correlated with increased Notch signaling ^27,28^. Therefore, expression of Notch signaling genes was investigated in the spheroids. As shown in Figure 10A and B, there was an increase in Notch1 and Notch3, Jag2 and Dll4 and the downstream genes Hes1 and Heyl, as well as, FSH receptor on spheroid formation. Thus, an up-regulation of Notch signaling correlates with the increase in FSHβ and FSHR expression in the ovarian cancer spheroids. Further, as shown in the Figure 11A, the mean size of the aggregates formed was significantly less in the presence of RF5 a/s and ScFv42, and this effect was significantly enhanced in presence of both inhibitors.

**Figure 10.**
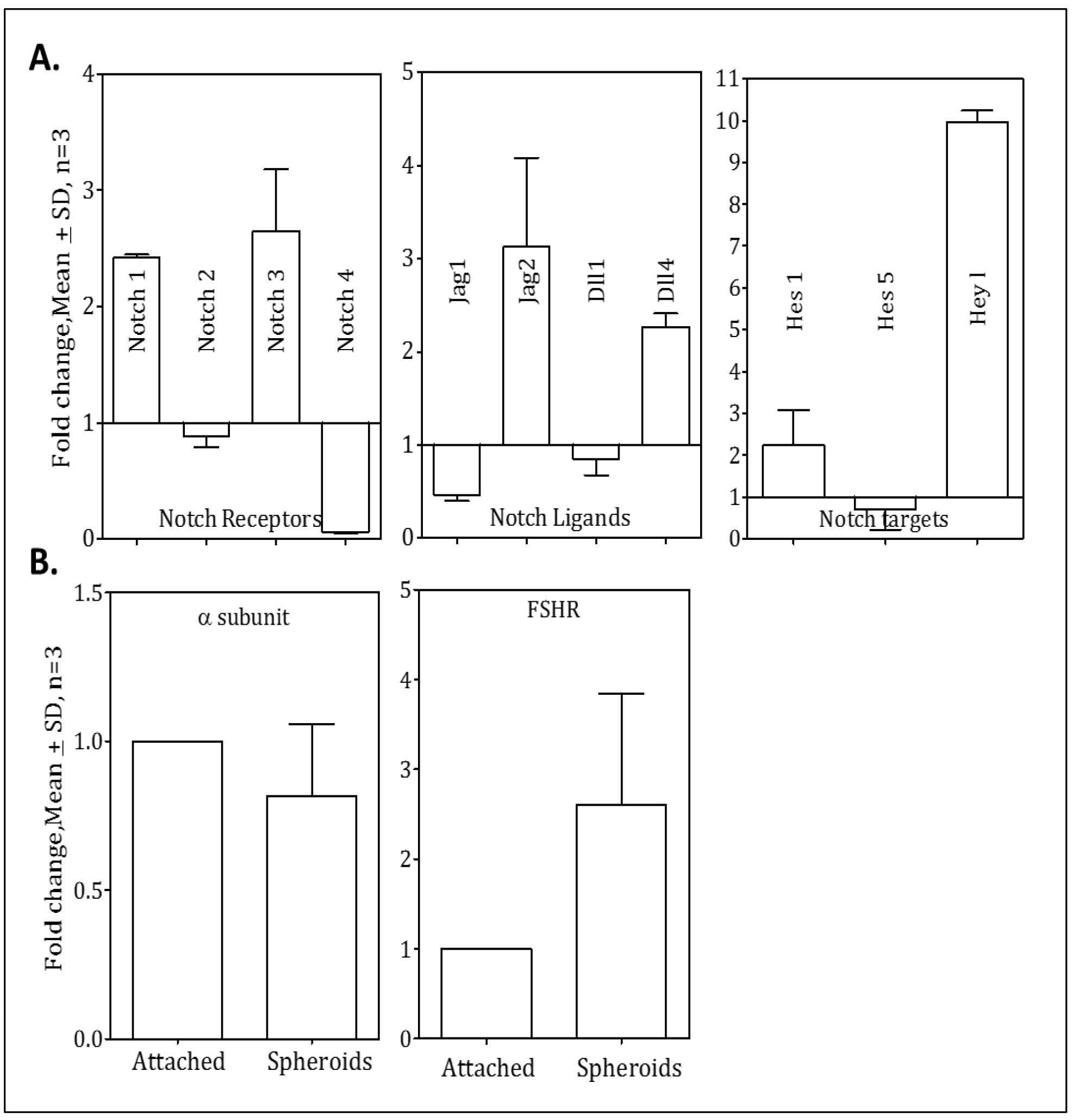
Status of Notch and FSH signaling genes in OVCAR-3 spheroids. The OVCAR-3 cells were cultured in low attachment conditions or as monolayer and harvested after 48 hours. Real time PCR was performed to assay the transcript levels of indicated Notch receptors, ligands targets and **B**. FSHR and α subunit. The fold change in the expression level of genes was calculated with respect to OVCAR-3 cell cultured as monolayer after normalizing with GAPDH expression. Error bars represent mean ± SD, n is the number of repeats.

**Figure 11.**
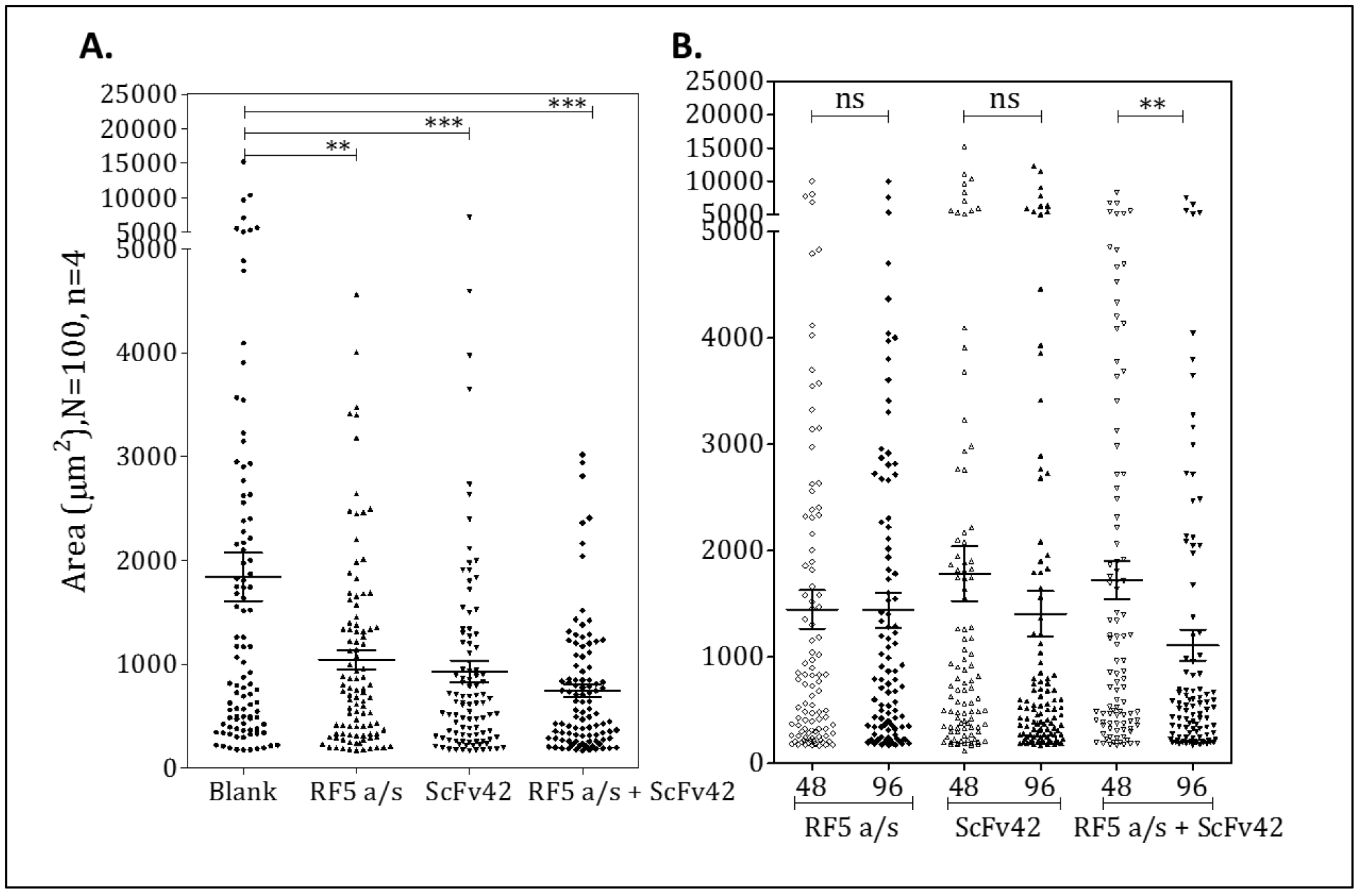
Effect of FSH and Notch antagonist (RF5 a/s and ScFv42 respectively) on the aggregation of OVCAR-3 spheroids. **A.** The OVCAR-3 cells were cultured in low attachment conditions in the presence of RF5 a/s (5μg/ml) and ScFv42 (10μg/ml) individually and in combination for 48 hours and the images were taken (5 fields/well) before and after the treatment. The area of the spheroids in each image was analyzed in MATLAB. The distribution of area of the 100 OVCAR-3 spheroids cultured in the presence and absence of antagonists is plotted. Each symbol represents the area of single spheroid. **B.** OVCAR-3 spheroids cultured for 48 hours were incubated with either RF5 a/s (5μg/ml), and/ or ScFv42 (10μg/ml) and the microscopic images were taken at regular intervals as above. The area of the spheroids in each image was analyzed in MATLAB. Distribution of area of the 100 spheroids was measured at 48 hours (before the addition of antagonists) and at 96 hours (i.e. two days after addition). Each symbol represents one spheroid and the solid symbols represents the spheroids incubated with different antagonists. Significance is calculated by comparison of treated and basal values using ANNOVA. Error bars represent mean± SEM. n represents the number of repeats and N is the number of spheroids analyzed. ns represents P> 0.05, * represents P≤ 0.05, ** represents P≤ 0.01 and *** represents P≤ 0.001 compared to non-treated.

In the next experiment, effect of both these inhibitors on the preformed spheroids was investigated. The OVCAR-3 cells aggregates were formed for 48 hours and then either RF5 a/s or ScFv42 individually, or in combination were added. As shown in Figure11B, there was a decrease in the mean aggregate size when both FSH and Notch signaling were inhibited.

## Discussion

The etiology of ovarian carcinomatosis is not very well understood. Both FSH and Notch have been shown to be involved in ovarian tumor angiogenesis, epithelial to mesenchymal transition, resistance to chemotherapy and cancer stem cell activity ^2,10^. Moreover, FSH has been shown to elevate Notch1 expression by the hormone in SKOV-3 cells ^29^. This study demonstrates a clear association between FSH and Notch in the context of ovarian cancer metastasis. Evidence for active FSH signaling in the three-distinct ovarian cancer cell lines has been demonstrated. All three cell lines express FSHR with affinities in the physiological range with the OVCAR-3 cell line showing highest FSHR density, as well as the response to the hormone. Presence of FSHR in cell lines was also confirmed by specific binding of FSHR-ECD a/s, which inhibited hormone binding (unpublished data), as well as response indicating the antagonistic nature of this a/s.

FSH increased overall Notch signaling by upregulation of different Notch receptors, ligands and the down-stream target genes and in presence of both FSH and Dll4, there was an additive increase in the Notch signaling. Increased proliferation of the ovarian cancer cell lines by FSH, which agrees with previous reports ^6,30^ was mediated by increased Notch signaling as it was inhibited by Notch3 NRR-specific ScFv. The proliferation was brought even below the basal rates by the combination of Notch3 NRR ScFv and FSHR a/s.

Progression of ovarian cancer frequently leads to accumulation of ascites in the peritoneal cavity of the patients which provides a complex mixture of soluble factors and cellular components creating a pro-inflammatory and tumor-promoting microenvironment for the cells ^31^. FSH is probably one such factor and its levels in the ascites of twelve ovarian cancer patients out of twenty-six patients were significantly high. Earlier reports ^32^ also indicated the serum/tumor fluid FSH ratios to be low in the ovarian cancer patients suggesting that the origin of FSH in the ascites may not be the serum, which could be higher in the post-menopausal status of many of the patients ^33^. The present study is however the first to posit that the spheroids obtained from the ascites are the source of FSH. We observed that OVCAR-3 and SKOV-3 cells expressed FSHβ mRNA and secreted FSH into the medium only when cultured as spheroids, and not when grown as monolayers. Expression of the α subunit mRNA was observed in the cancer cell lines cultured both as monolayers and as spheroids, as well as, the spheroids obtained from the patients. In contrast, FSHβ expression is seen only when the cells are organized into the spheroidal geometries. In support of our cell line-based studies, the expression of FSHβ (and even FSHα) was observed to be very low in cells from the primary ovarian patient tumor samples. Despite low ligand expression, FSHR expression was high. This suggests that FSH secreted by the ascitic spheroids might not just impact the morphogenesis of the latter, but also turn on FSH signaling in the FSHR expressing primary tumor driving its proliferation. This feedback loop, however, is dependent upon the establishment of the metastatic niche, and might be perhaps one of the earliest examples of how the metastatic niche feeds back to alter the behavior of the primary tumor niche of a cancer. Moreover, high levels of FSH in the ascites of the ovarian cancer patients are inversely correlated to their survival ^33^.

It is well established that FSHβ expression is the rate limiting step in the secretion of the hormone from the pituitary ^34^ and once the β subunit expression is turned ‘on’, the heterodimeric hormone is secreted into the environment. Turning ‘on’ of FSHβ expression in the spheroids is very intriguing and has not been reported earlier. Spheroids have been known to show altered epigenetic regulation with a general increase in histone acetylation ^35^. However, the effect on FSHβ expression appears to be specific as neither the expression of LHβ is upregulated nor was there any secretion of LH under these conditions. Therefore, whether such a spheroid-specific expression and secretion of FSH is prognosticative of a metastatic, niche-specific secretome needs to be investigated further.

It is evident that FSH expressed by the ovarian cancer spheroids upregulates Notch signaling, which together with the stromal Dll4 leads to increased proliferation of the tumor cells. It was also interesting that the spheroidal cells also exhibited higher expression of FSHR probably making them extremely sensitive to the increasing concentrations of the hormone. These data indicate that there is an autocrine regulation of FSH-FSHR signaling that plays an important role in the ovarian cancer progression.

The ovarian cancer cell spheroids are known to be associated strongly with the recurrence of the ovarian cancer, as they show resistance to chemo- and radio-therapies with larger spheroids being more resistant to the therapies ^13^. Under these conditions, both FSH and Notch pathways are interesting immunotherapeutic targets. The Notch NRR ScFv42, individually, and in combination with FSHR a/s significantly inhibited formation of bigger OVCAR-3 spheroids. The antibodies also decreased the average size of the pre-formed spheroids. These two antibodies also inhibited ovarian cancer cell proliferation, further suggesting the therapeutic potential of these antibodies. Our observation that the FSHR antibody can perturb spheroidal morphogenesis individually but inhibit proliferation of monolayers only in an FSH-added condition suggests a morphology-specific higher activity of FSH signaling and its integration with the Notch signal transduction. The mechanistic investigations of such signaling crosstalk will be explored in future endeavors.

In conclusion, this study establishes a clear link between FSH and Notch in the ovarian cancer progression and show that these are the potential targets for future therapeutic efforts.

## Additional Information

### Authors’ contributions

SG, RB^1^ and RRD designed the study and wrote the manuscript. SG carried out the execution of the project. SKS, SNS, RB^2^, AV and RG were involved in identifying and characterizing the ovarian cancer patients. RB^1^: Ramray Bhat; RB^2^: Rahul Bhagat

### Ethics approval and consent to participate

The studies described here were carried out as per the guidelines of the Institutional review board and in agreement with the ethical guidelines of the Kidwai Cancer Institute, Sri Shankara Cancer Hospital and Research Centre, and the Indian Institute of Science (IISc) after obtaining the informed consent from the patients.

## Acknowledgements

The authors thanks Karthik Murthy and Amith Kumar (Center for Ecological science, Indian Institute of Science) for developing MATLAB script for the analysis of area under spheroids, and Purba Sarkar and Shyamili Goutham for help with primary tumor collection, RNA isolation and cDNA synthesis.

